# Sex-dependent additive effects of dorzagliatin and incretin on insulin secretion in a novel mouse model of *GCK*-MODY

**DOI:** 10.1101/2024.11.09.622781

**Authors:** Shadai Salazar, Luis Fernando Delgadillo-Silva, Priscila Carapeto, Karen Dakessian, Rana Melhem, Audrey Provencher-Girard, Giada Ostinelli, Julie Turgeon, Imane Kaci, Francis Migneault, Mark O. Huising, Marie-Josée Hébert, Guy A. Rutter

## Abstract

Glucokinase (GK) catalyses the key regulatory step in glucose-stimulated insulin secretion. Correspondingly, hetero– and homozygous mutations in human *GCK* cause maturity-onset diabetes of the young (GCK-MODY) and permanent neonatal diabetes (PNDM), respectively. To explore the possible utility of glucokinase activators (GKA) and of glucagon–like receptor-1 (GLP-1) agonists in these diseases, we have developed a novel hypomorphic *Gck* allele in mice encoding an aberrantly spliced mRNA deleted for exons 2 and 3. In islets from homozygous knock-in (Gck^KI/KI^) mice, GK immunoreactivity was reduced by >85%, and glucose-stimulated insulin secretion eliminated. Homozygous Gck^KI/KI^ mice were smaller than wildtype littermates and displayed frank diabetes (fasting blood glucose >18 mmol/L; HbA1c ∼12%), ketosis and nephropathy. Heterozygous Gck^KI/+^ mice were glucose intolerant (HbA1c ∼5.5%). Abnormal glucose-stimulated Ca^2+^ dynamics and beta cell-beta cell connectivity in Gck^KI/+^ islets were completely reversed by the recently-developed GKA, dorzagliatin, which was largely inactive in homozygous Gck^KI/KI^ mouse islets. The GLP-1 receptor agonist exendin-4 improved glucose tolerance in male Gck^KI/+^ mice, an action potentiated by dorzagliatin, in male but not female mice. Sex-dependent additive effects of these agents were also observed on insulin secretion *in vitro*. Combined treatment with GKA and incretin may thus be useful in *GCK*-MODY or *GCK*-PNDM.

**Article Highlights:**

a. Glucokinase deficiency can drive maturity-onset diabetes of the young (GCK-MODY; *heterozygotes*) and permanent neonatal diabetes (GCK-PNDM; *homozygotes*)
b. We describe a hypomorphic *Gck* allele where aberrant splicing in islets lowers GK activity to by ∼85%. We use these mice to explore the effects of the glucokinase activator, dorzagliatin, and incretin on insulin secretion
c. Whereas heterozygous mutant mice are mildly hyperglycemic, homozygotes have frank diabetes but survive to adulthood. Dorzagliatin potentiates the effects of GLP-1 receptor activation sex-dependently in heterozygotes
d. Combined use of these drugs may be useful in some forms of *GCK* diabetes

## Introduction

Diabetes affects more than 1 in 10 of the global adult population (1), a figure expected to grow to >750m people by 2045 (2). Glucokinase (Hexokinase-4, HK-IV, EC 2.7.1.1, GK) catalyses the flux-generating step in glycolysis and has been dubbed the “glucose sensor” of the pancreatic beta cell (3–7). After phosphorylation by GK, glucose carbons flow through the glycolytic pathway and enter the citrate cycle, stimulating respiratory chain activity to increase cytosolic ATP/ADP ratios. Closure of ATP-sensitive K^+^ channels leads to depolarisation of the plasma membrane, Ca^2+^ influx through voltage-gated calcium channels, and insulin release (8; 9). Roles for other coupling factors (10; 11), and local ATP/ADP microdomains (12), are also proposed (13), though the latter are disputed (14).

Homozygosity for GK (*GCK)* loss-of-function (LoF) alleles results in permanent neonatal diabetes mellitus (PNDM) (15), often requiring insulin treatment shortly after birth (16–18). Heterozygosity for LoF mutations is associated with maturity onset diabetes of the young (MODY) (19; 20). *GCK*-MODY (formerly MODY2) accounts for 20-30% of all MODY cases. Although chronically hyperglycemic (fasting glucose 6-8 mM) (21), *GCK*-MODY patients are largely asymptomatic (21). However, fetal macrosomia is an important complication in pregnancy (22).

Several mouse models have been generated to explore how *Gck* insufficiency impacts whole body metabolism (23). Animals deleted for exon 2 and incorporating a frameshift mutation (24), or exon 4 plus parts of exons 3 and 5 (25), respectively show *in utero* or perinatal mortality as homozygotes, and mild hyperglycemia as heterozygotes. Beta cell-selective *Gck* knock-outs display severe hyperglycemia, and die a few days after birth (26; 27). Demonstrating the importance of islet dysfunction in the impact of these mutations, rescue of *Gck* expression selectively in the beta cell of *Gck* null mice is sufficient to reverse lethality (25).

Incretin drugs including glucagon-like-1 (GLP-1) receptor agonists are now used widely in the clinic and provide a highly efficient treatment for T2D (28). Use of these agonists has not, up to now, been explored either in *GCK*-MODY or *GCK*-PNDM. In any case, incretin action on insulin secretion requires glucose metabolism (28), suggesting that the effects of these agents may be limited in the above diseases.

Dorzagliatin (29) is a latest-generation allosteric glucokinase activator (GKA), that has recently been shown to be therapeutically useful in *GCK*-MODY (30). The above findings raise the possibility that the combined use of incretins and GKA may provide additional benefits compared to the use of either agent alone.

Here, we explore this possibility in wildtype mice and in animals bearing a hypomorphic *Gck* allele that results in aberrant splicing and severe depletion of active GK protein. Residual GK activity is nevertheless sufficient to allow the survival of homozygous mice into adulthood, despite severe diabetes, modelling *GCK*-PNDM. Heterozygous Gck^KI/+^ mice provide a convenient model of *GCK*-MODY. We test the potential therapeutic utility of glucokinase activators and incretins in both models.

## Materials and Methods

### Generation of hypomorphic *Gck* (C57BL6/J-Gck^tm1(mCard)/Rutt^, Gck^KI^) alleles

The overall strategy, as designed with GenOway (Grenoble, Fr), is illustrated in Fig. 1A and SFig. 1. Integration of the mutant allele was confirmed by Sanger sequencing. Assessed after mRNA extraction from isolated islets of homozygous male Gck-mCardinal (*Gck*^KI/KI^) mice, amplification by conventional PCR of the region from exon 1 to exon 9 of mRNA-derived cDNA from (SFig. 1B) yielded a barely-detectable band corresponding to the predicted wild-type (WT) product (∼1.17kb) alongside a major product of ∼850bp (∼350bp less than predicted). Sanger sequencing of the latter revealed an aberrant splicing event between exons 1 and 4, eliminating the whole of exons 2 and 3 (Fig. 1B, SFig. 1C). *In silico* translation of the aberrantly-spliced isoform suggested the production of a 360 amino acids (AA) protein versus native wild type (466 AA) glucokinase. The mutant form is expected to lack amino acids 16 to 121, including residues in the small lobe encoded by exon 3, and critical for ATP binding and catalysis, e.g. R63 (6; 31; 32). cDNAs corresponding to the expected IRES or mCardinal regions were not detected in islets, and mCardinal fluorescence was absent from islet, liver and brain (ventromedial and lateral hypothalamus) (33) (not shown).

**Figure 1.**
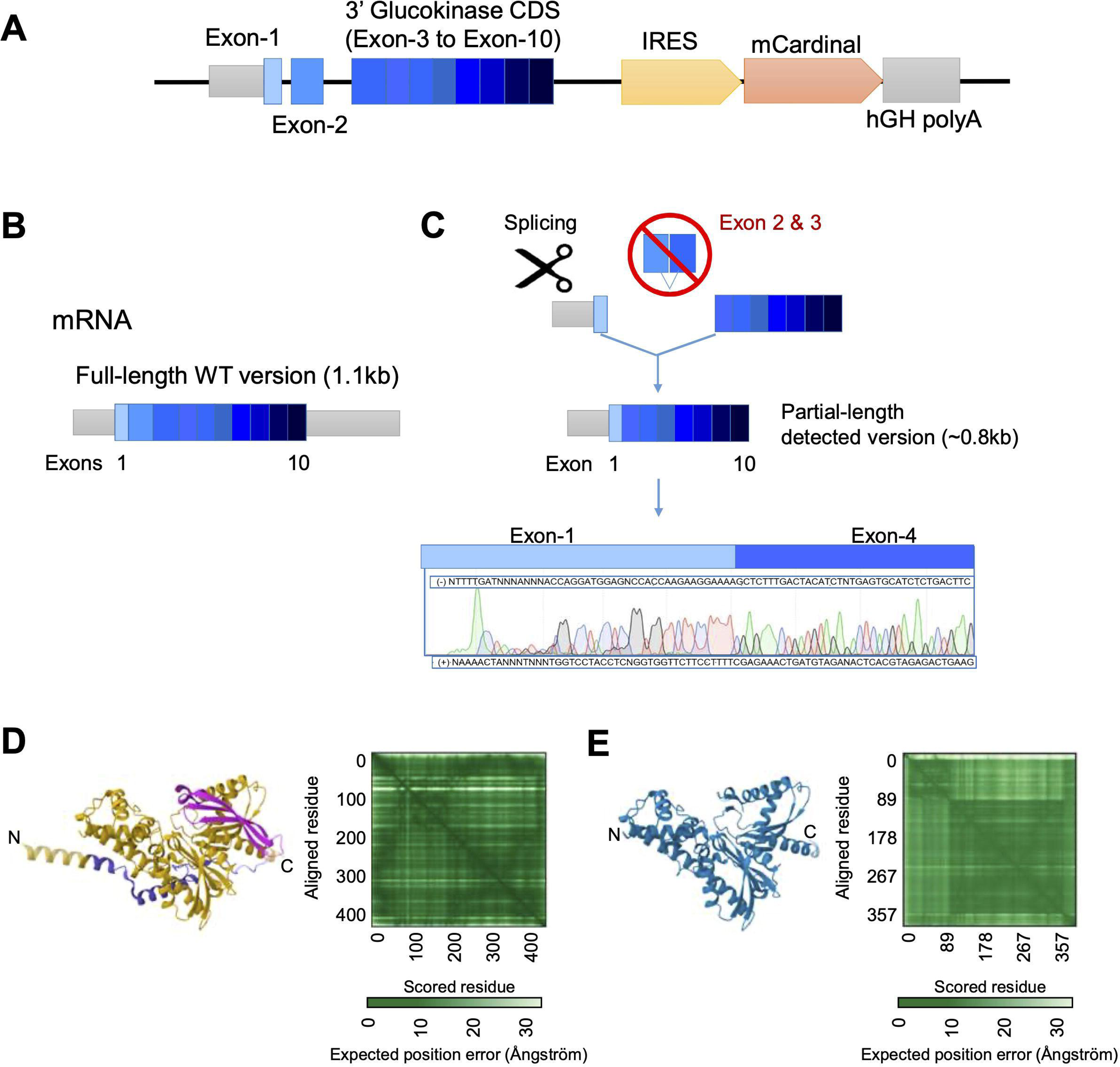
Design and genomic map of the Gck^KI^ mouse allele. **A)** The 3’ *Glucokinase* (*Gck*) coding sequence comprising exon-4 to exon-10 was directly fused in frame with endogenous exon-3. This was followed by an IRES, mCardinal cDNA and termination signal plus human growth hormone (hGH) polyA. The construct is expected to produce a single bicistronic mRNA expressing Glucokinase and mCardinal from the same mRNA transcript. **B)** Wildtype *Gck* mRNA structure. **C)** Observed sequence expressed from the knock-in allele. Sanger sequencing of the *Gck* PCR product from Gck^KI/KI^ islet cDNA revealed alternative splicing in which exon 2 and exon 3 are skipped. Alpha-fold predicted 3D structure of **D)** wildtype mouse glucokinase and **E)** the mis-spliced isoform. The green matrix represents the confidence in the prediction in Angstroms, with darker shades indicating greater confidence. The beta-barrel that forms part of the glucose-binding pocket and ATP binding domain (pink, D) is absent from the modelled mis-spliced isoform (E).

### Animal husbandry

All experimental manipulations were approved by the local ethical committee (CRCHUM, Montreal CIPA 2022–10,040 CM21022GRs). Colonies of GK^KI^ and of GK^KI^:Ins1Cre:GCaMP6^f/f^ and GK^KI^:Ins1Cre:GCCaMP6^f/f^ mice, in which the Ca^2+^ sensor GCaMP6f is expressed selectively in the beta cell (34), on a C57BL/6J background, were fed a regular chow diet and maintained at controlled temperature (21–23 °C), humidity (45–50 %) and light (12 h day-night cycle).

### Structural modelling

We compared the Alpha fold-predicted 3D protein structure against the X-ray crystal-resolved human Glucokinase (3VEY, www.rcsb.org/structure/3VEY). Additionally, we compared the previously AI-generated model of mouse Glucokinase (35). The 3D comparison was generated by PDB Mol View (Pairwise Structure Alignment https://www.rcsb.org/alignment) using the TM-algin method. The TM-align method is based on structure comparison for proteins with similar global topology. ClustalW was employed to quantify amino acid sequence similarity.

### Intraperitoneal glucose tolerance tests (IPGTT)

Mice were fasted overnight (16 h) with free access to water. At 9 am, glucose (2 g/kg body weight) was administered *via* intraperitoneal injection. Blood glucose levels were measured from the tail vein 0, 15, 30, 60 and 90 min. later with an automatic glucometer (Contour next ONE; Canada) (36).

### Insulin tolerance tests

After 6 h fasting with free access to water, human insulin (Novolin-ge Toronto, Novo Nordisk) (1 U/kg body weight) was administered *via* intraperitoneal injection. Blood glucose was measured as above (36).

### Islet isolation

Islets were isolated from mice aged between 8 and 16 weeks, essentially as previously described (37).

### Measurement of insulin secretion *in vitro*

Initially, insulin secretion was assessed using the following buffer, containing (mmol/L) 137 NaCl, 4.8 KCl, 1.2 KH_2_PO_4_, 1.2 MgSO_4_, 2.5 CaCl_2_·2H2O, 5 NaHCO_3_, and 16 HEPES, adjusted to pH 7.4 and supplemented with 0.1% BSA. Islets were pre-incubated in low glucose (3 mM) for 1 h, the sequentially for 1 h at 3 mmol/L then 17 mmol/L glucose (see Fig. 3, H,I). In experiments involving agonist treatment (Fig. 6), isolated islets were incubated in modified KRBH stock solutions comprising (mmol/L) 10 HEPES, 2 NaHCO_3_, 137 NaCl, 3.6 KCl, 0.5 NaH_2_PO4, 0.5 MgSO4, 1.5 CaCl_2_. Solutions were gassed with 95% O_2_ and 5% CO_2_ and adjusted to pH 7.4. The buffer was maintained at 37°C and supplemented with 0.1% BSA. Islets were selected based on size, washed, and incubated in KRBH-BSA with 3 mmol/L glucose for 1 h then transferred to 12-well plates and divided into four groups, each in triplicate: (1) control (2) plus Dorzagliatin (10 µM), (3) Exendin-4 (100 nmol/L), (4) Dorzagliatin and Exendin-4. Islets were incubated for 30 min. in low glucose (3 mmol/L), then at high (17 mmol/L) glucose. All samples were stored at –80°C until measurement using an Insulin Ultrasensitive HTRF Assay Kit (Revvity, 62IN2PEG).

### Beta– and alpha-cell mass

After transcardiac perfusion, the pancreas was carefully dissected, weighed, and placed in cassettes for fixation in 4%(v/v) paraformaldehyde (PFA) for 2h to facilitate paraffin embedding. Adjacent full footprint sections were then subjected to immunostaining for glucagon (Cell Signaling, 2760) or insulin (Cell Signaling, 4590) using a standard DAB substrate kit (Cell Signaling, 8059). Incubations with primary antibodies were performed overnight at 4°C, at an antibody dilution of 1:100. The mass of β-cells and α-cells was quantified using point counting morphometry (38).

### Renal histology

Whole kidney was dissected after transcardiac perfusion and fixed in 4%(v/v) PFA for an extra 2 h, then paraffin-embedded. Histology analysis was done at a 20x magnification, on whole kidney sections. Images were taken with Slide Scanner Leica, Aperio Versa 200 digital scanner and further analyzed in the Aperio ImageScope 12.43.3 software. Renal tubular damage was described based on luminal dilatation and necrosis, loss of brush border, and cast formation. The investigator was blinded to experimental conditions.

### PCR and qPCR

Specific primers for *Glucokinase* cDNA amplification from exon 1 to the mCardinal sequence are reported in Supp. Table 1. Briefly, was cDNA synthesized from total Trizol-Chloroform purified mRNA from isolated islets of either Gck^KI/KI^ or WT animals. cDNA was synthesized using High-Capacity cDNA Reverse Transcription kit (Applied biosystems, Thermo-Fisher). After PCR amplification of exon 1 and exon 9, the product was migrated on a 1.0% agarose gel, the ∼850bp band was excised, purified and Sanger sequenced (Genome-Quebec, Canada).

### Calcium imaging

Ca^2+^ imaging was performed essentially as described (39; 40) using a Zeiss LSM 900 Airyscan 2 super-resolution confocal microscope, equipped with an incubation system set at 37°C. A 40×/1.3 Apochromat oil immersion objective was employed with a frame rate of 150ms (∼6Hz) with 512 x 512 px image size. 1h before imaging, islets were transferred into Krebs buffer (130 mM NaCl, 3.6 mM KCl, 1.5 mM CaCl_2_, 0.5 mM MgSO_4_, 0.5 mM NaH_2_PO_4_, 24 mM NaHCO_3_, 10 mM HEPES; pH 7.4), containing 3 mmol/L glucose. For Dorzagliatin (HMS5552; Abmol Biosciences) (41; 42) treatment, islets were transferred to Krebs buffer with 1% DMSO and 10 µm Dorzagliatin for 1h prior to imaging. Imaging was performed at 11 mM glucose.

### Connectivity analysis

Analysis was performed essentially as per (39), with modifications. Fluorescence traces were normalized using the formula:

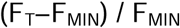

Smoothing was used to adjust Ca^2+^ signals via a moving average filter, contributing to 1% of the total length of the Ca^2+^ recording. Cell activity was represented in binary form, where any time point deviating >20% above the baseline was considered to be active, represented by “1”. Any inactive time point, under the threshold, is represented by a “0”. The following formula was used to calculate coactivity for each cell pair:

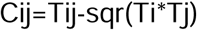

*Tij* denotes the total coactivity of each cell pair. *Ti* and *Tj* are the total activity time for the two cells compared, respectively.

Cells are considered to be linked if their t-test displayed a chance (>2 standard deviations i.e. p<0.01) probability within a corresponding distribution of the shuffled (10,000 times) binarized observed dataset, for the analyzed cell pairs.

A topographic representation of the connectivity was plotted in MATLAB (available from: https://www.mathworks.com-matlabcentral-fileexchange-24035-wgplot-weighted-graph-plot-a-better-version-of-gplot). Edge colors indicate the strength of the coactivity. Highly connected (“Hub”) cells were defined as those where >70 % of connections had a connectivity value of >0.7.

### Western (immuno-) blotting

Blotting was performed *as per* (43). Isolated islets were washed twice with PBS, and fast frozen on dry ice. To extract proteins, 10uL of RIPA buffer plus protease inhibitors was added to 100 islets, and samples sonicated for 3s on ice. Protein concentration was estimated by Pierce BCA protein assay (ThermoFisher). One volume of Laemmli (4x) solubilization buffer was added, followed by exposure to 95LC for 60s to denature proteins. 10µg protein per lane was loaded onto 10%(w/v) acrylamide gels. For liver, lung and heart, 10uL RIPA buffer with protease inhibitors per mg wet weight was added. Protein extract (20µg) was loaded onto each lane. After electrophoresis, proteins were transferred to PVDF membranes. Membranes were blocked with 3% BSA, TBST 1%. After 1h, membranes were incubated with primary antibody overnight at 4LC. Membranes were washed with TBST 1% before incubation with secondary antibody (1h, room temperature). Horseradish peroxidase (HRP) chemiluminescent substrate (ECL western substrate, Thermo) was used to reveal bound antibody.

### Statistics

Data are expressed as mean ± SD, unless otherwise stated and significance tested by one or two-way ANOVA with Šidák or Brown-Forsythe multiple comparison tests, using GraphPad Prism 9 (GraphPad Software, San Diego, CA). *P* < 0.05 was considered significant.

## Data resource and availability

All data generated or analyzed during this study are included in the published article (and its online supplementary files. Details of movies are provided in the Supp. Figure document, which includes original gel blots in SFig.s 8,9). Analytical scripts are provided at https://zenodo.org/records/14042795.

## Results

### Generation of a hypomorphic Gck allele

With the initial objective of identifying beta cell subpopulations enriched for GK (which may correspond to highly connected “hub” cells) (44), we designed a “knock-in” construct encoding exons 3-10 of mouse *Gck*, followed by an internal ribosome entry site (IRES), cDNA encoding the fluorescent protein mCardinal, and a polyA sequence (Materials and Methods, Fig. 1A,C, SFig. 1A). Genomic sequencing confirmed the expected integration at the *Gck* locus (not shown). However, and unexpectedly, an alternatively-spliced product was generated, predicted to encode a mutant protein that lacks much of the small lobe of GK (31).

GK protein levels in islets from wild type and mutant mice were assessed by Western (immuno-) blotting (Fig. 2). *Gck* is expressed from different promoters in the liver and in the pancreatic beta cell (45). Consequently, the first 15 amino acids (AAs) encoded by exon 1 from the beta cell promoter differ from those of the equivalent exon encoded in the liver (6). The antibody deployed (Thermofisher Glucokinase Antibody PA5-15072) is raised against the first 30 AA of type 1 GK (islet). Loss of the following 16 AA (encoded by exon 2) from the epitope is thus likely to weaken or destroy recognition by the antibody of the mis-spliced isoform. This approach revealed a 30-45% and ≥85% lowering in apparent GK immunoreactivity (∼48 kDa) in islets from GK^KI/+^ and GK^KI/KI^ islets, respectively, *versus* control mice of either sex, indicating a proportionate lowering of the intact isoform 1 protein (including exons 2 and 3) (Figure 2B). We note that, if present, the mutant is expected to be inactive, given (a) the requirement for the ATP binding sites encoded by exon 3 (6) and (b) the likely absence, based on structural prediction with AlphaFold (see Methods, Fig. 1D,E, S.Movie1), of a properly-folded glucose-binding beta sheet in the small lobe.

**Figure 2.**
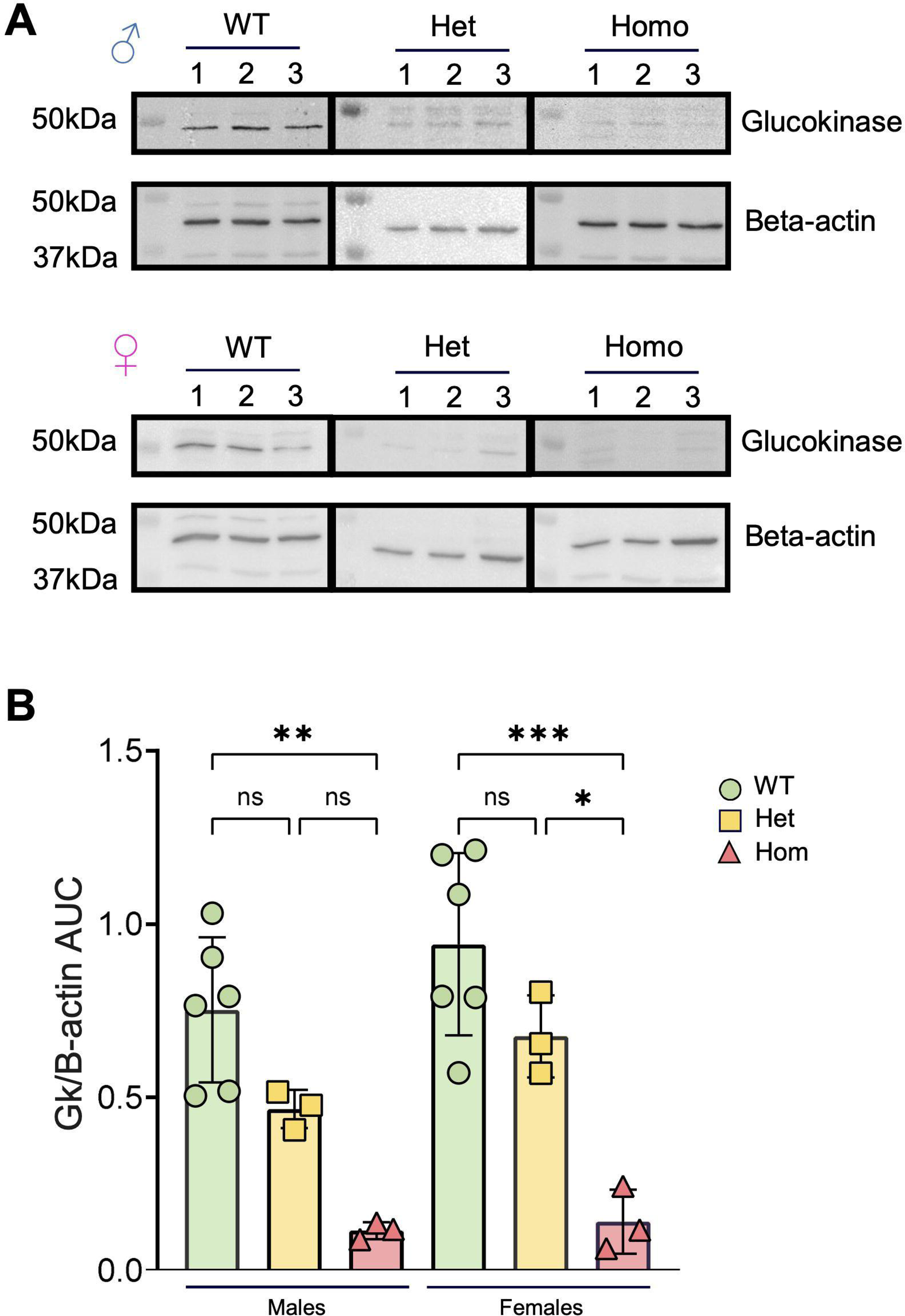
Apparent GK protein levels in male and female pancreatic islets. **A)** Western (immuno-) blot images from islets protein from wildtype, Gck^KI/+^ (heterozygous) and Gck^KI/KI^ (homozygous) males and females. Glucokinase was detected at ∼48kDa and beta-actin at ∼45kDa. **B)** Quantitative analysis of GK expression levels normalized to beta-actin across sex and genotype. Each dot represents a single animal. Unpaired one-way ANOVA with Tukey’s correction. \**P* ≤ .05, \*\**P* ≤ .01, \*\*\**P* ≤ .001.

No differences in immunoreactivity at this molecular weight were observed in liver extracts from WT, Gck^KI/+^ and Gck^KI/KI^ animals, presumably reflecting the failure of the antibody to recognise the liver isoform (SFig. 2).

### Metabolic characterisation of *Gck* mutant mice

Examined at 8 weeks of age, heterozygous (Gck^KI/+^) mice displayed normal body weight, whilst male homozygous (Gck^KI/KI^) animals were lighter than littermate controls (Fig. 3A,B); a similar tendency was seen in females (Fig. 3B). HbA1c levels tended to be elevated, or were drastically increased, respectively, in Gck^KI/+^ and Gck^KI/KI^ animals *versus* controls (Fig. 3C). Fasting blood glucose (FBG) was >18 mmol/L in the latter animals of either sex at eight weeks of age (males, *n*=7, 21.6 to>33.3 mmol/L; females, *n*=5, 18.4-27.4 mmol/L; Fig. 3D,F). Whereas overnight (16 h) FBG did not differ between Gck^+/+^ and Gck^KI/+^ mice of either sex (Fig. 3D,E), Gck^KI/+^ animals displayed markedly abnormal glucose tolerance, with no return to pre-infusion levels after 90 min. Blood glucose levels after 6 h fasting were higher in Gck^KI/+^ *versus* WT mice (Fig. 3F,G,). Insulin sensitivity did not differ between genotypes, after correction for FBG (Fig. 3F *insets*, Fig. 3G).

**Figure 3.**
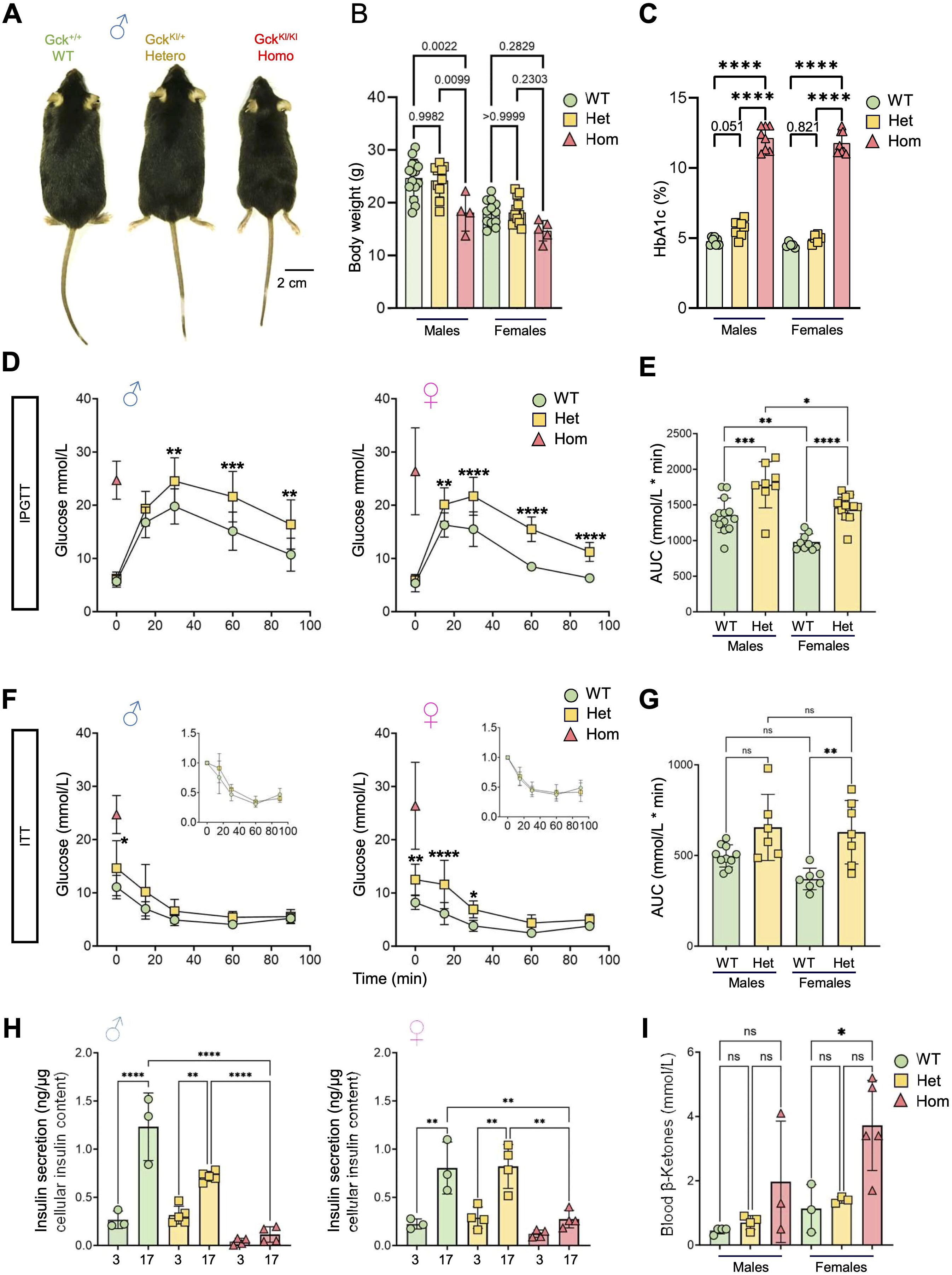
Metabolic characterization of Gck^KI/+^ and Gck^KI/KI^ mice. **A)** Representative images across male genotypes. **B)** Mouse weight at 8 weeks of age across sex and genotype, *n*=15 WT, 10 Het and 4 Homo. Females, *n*=13 WT, 13 Het and 5 Homo. **C)** HbA1c at 8-16w across sex and genotype. Homozygous animals exhibit a diabetic phenotype in both males and females (HbA1c% > 10). Males, *n*=7 WT, 8 Het, and 8 Homo. Females, *n*=5 WT, 6 Hets and 8 Homo. Unpaired one-way ANOVA with Tukey’s correction. **D)** Intraperitoneal glucose tolerance tests (IPGTT) with 2 g/kg glucose performed at 8-16w of age for males (*n*=12 WT and 9 Hets) and females (*n*=12 WT and 13 Hets). Unpaired one-way ANOVA with Šidák’s correction. **E)** Glucose excursion (area under the curve, AUC) quantifications from the male and female IPGTTs presented in (D). Unpaired one-way ANOVA with Tukey’s correction. **F)** Intraperitoneal insulin tolerance test (IPTT) with 0.75 insulin units/Kg performed at 8-16w. Males, *n*=10 WT, 6 Hets. Females, *n*=7 WT, 7 Hets. Unpaired two-way ANOVA with Šidák’s correction. **G)** Glucose AUC quantifications of IPGTTs presented in (F). Unpaired one-way ANOVA with Tukey’s correction. The homozygous fasting values are represented by the red triangle (*n*=3 mice) in (D) and (F). Due to the high fasting glucose, IPGTT and IPTT were not performed in homozygotes, and the fasting glucose levels are presented as a reference only. **H)** Insulin secretion from isolated islets across genotypes and sexes. * (I) Ketone body levels in fasted animals. *N*=3-5 per group. *P* ≤ .05, \*\**P* ≤ .01, \*\*\**P* ≤ .001, \*\*\*\**P* ≤ .0001.

Measured in batch incubations, glucose-stimulated insulin secretion (GSIS) from male islets was lowered by ∼50 % in Gck^KI/+^ versus controls, and eliminated in homozygous Gck^KI/KI^ islets (Fig. 3H). In females, GSIS was unaffected in Gck^KI/+^ but eliminated in Gck^KI/KI^ islets (Fig. 3H).

Whilst also unaltered in Gck^KI/+^ mice *versus* controls, beta cell mass was substantially (∼65% in males, ∼50 % in females) lowered in Gck^KI/KI^ animals (SFig. 3). Conversely, alpha cell numbers were markedly increased in Gck^KI/KI^ mice (∼2-fold in both sexes; SFig. 3). Consequently, the beta:alpha cell ratio fell from ∼10:1.0 in wildtype to ∼1.0:1.0 in Gck^KI/KI^ pancreata.

Consistent with frank diabetes, ketone levels tended to be, or were significantly, increased in male and female Gck^KI/KI^ animals *versus* controls, respectively (Fig. 3I). Moreover, in the kidney, homozygous Gck^KI/KI^ mice of either sex displayed clear tubular alterations compared to heterozygous Gck^KI/+^ and control mice (SFig. 4; Supp. Table 2).

**Figure 4.**
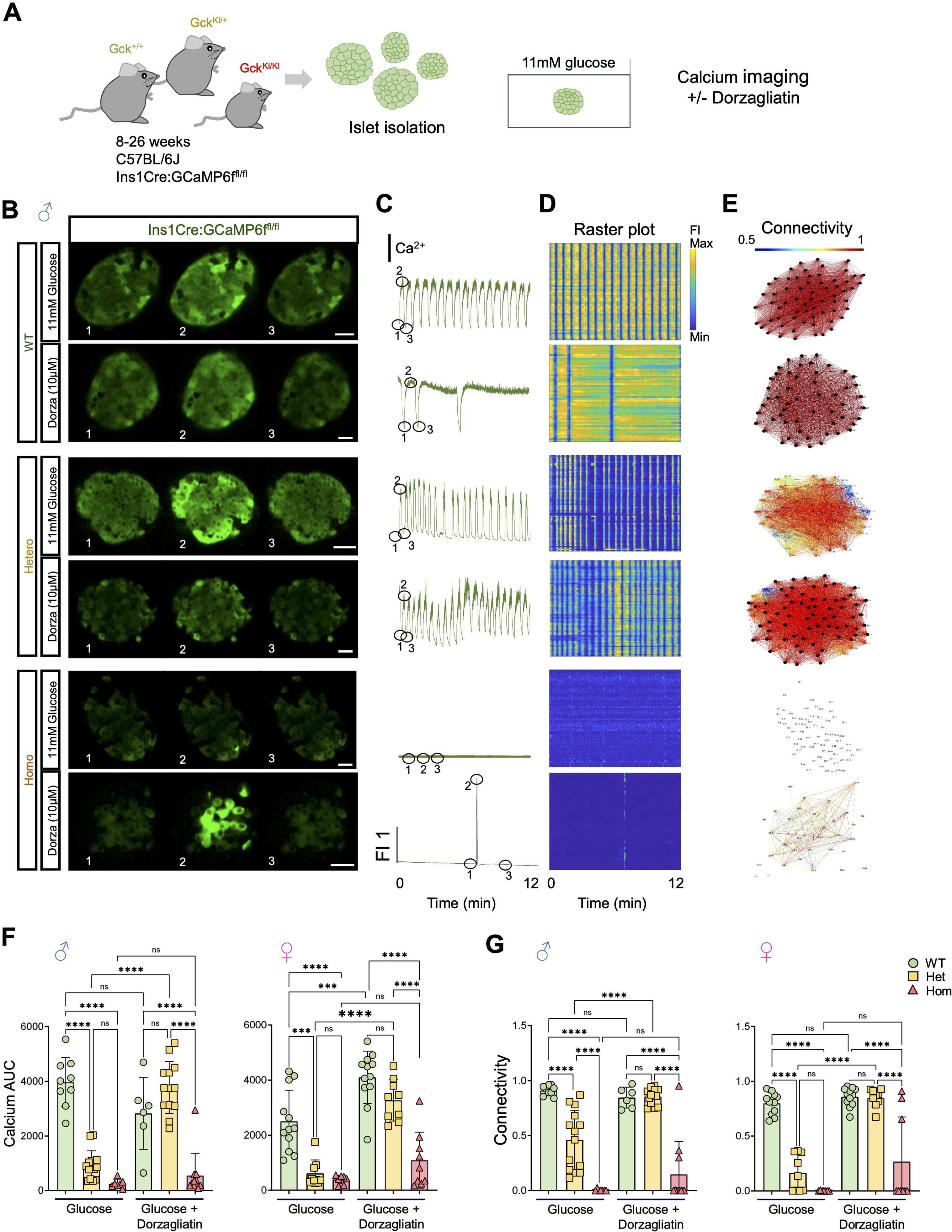
Dorzagliatin restores glucose-induced Ca^2+^ dynamics and β-cell connectivity in Gck^KI/+^ mouse islets. **A**) Experimental design for *in vitro* calcium imaging in islets from 8-16w Ins1Cre:GCaMP6f^fl/fl^:Gck^+/+^ (“WT”), Gck^KI/+^ (“Het”) and Gck^KI/+^ (“Homo”) triple transgenic mice. Where indicated, isolated islets were treated for 1h with 10µM dorzagliatin prior to imaging. **B)** Snapshots from confocal time-lapses recordings of the individual islets at 6 Hz in 11mM glucose with or without 10µmol/L dorzagliatin across *Gck* genotypes. **C)** Normalized GCaMP6f fluorescence traces from islets shown in (B). **C)** Raster plots show the apparent Ca^2+^ signal for individual cells from islets shown in (B). **D)** Visualization of the islet functional network for each genotype, with or without dorzagliatin, from islets displayed in (B). **E)** Calcium AUC quantification across genotypes in control and dorzagliatin treated islets. **F)** Connectivity quantifications across genotypes in control and dorzagliatin treatment groups. Unpaired one-way ANOVA with Tukey’s correction. \**P* ≤ .05, \*\**P* ≤ .01, \*\*\**P* ≤ .001, \*\*\*\**P* ≤ .0001. For 11 mmol/L glucose-only treatment, *n*=5 WT mice (9 islets), n=8 heterozygotes (14 islets) and n=4 homozygotes (7 islets). For dorzagliatin treatment, *n*=4 WT (6 islets), *n*=5 heterozygotes (13 islets) and *n*=4 homozygotes (11 islets). Each dot represents an individual islet. The numbers 1, 2, and 3 in the snapshots displayed in (B) indicate the time points corresponding to the fluorescence traces showed in (C). Scale bar, 25 μm.

### Ca^2+^ dynamics and intercellular connectivity

The above findings suggested that defects may exist in intracellular glucose handling or signalling in mutant beta cells. To explore this possibility, we studied glucose-regulated intracellular Ca^2+^ dynamics and cell-cell connectivity in isolated islets using high speed confocal imaging (Fig.4A) (44). To ensure that Ca^2+^ was measured exclusively in beta cells, mice were bred to animals carrying Ins1*Cre* alleles (46) and STOP-Flox alleles of the genetic Ca^2+^ sensor, GCaMP6f (34). Male animals carrying wild type *Gck* alleles displayed robust responses to stimulation with 11 mM glucose, and a high degree of connectivity (Fig. 4B-E; SMovie 2,3). These responses were significantly weakened in Gck^KI/+^ mice and almost completely eliminated in Gck^KI/KI^ animals. Similar behaviour was seen in islets from animals of both sexes, though the Ca^2+^ responses (area under the curve, AUC) were lower in females than males (Fig. 4, E-G SFig. 6). Remarkably, these responses were fully normalized with dorzagliatin in Gck^KI/+^ mice of both sexes Fig.4F,G, SFig. 6, SMovie 4,5), but barely affected in Gck^KI/KI^ mice. Nevertheless, a subset of female Gck^KI/KI^ islets showed detectable Ca^2+^ transients in response to glucose stimulation in the presence of dorzagliatin (Fig. 4, F,G).

### Responses of wildtype and Gck^KI/+^ mice to GKA and incretin *in vivo*

Given the findings above, we next explored the effects of dorzagliatin *in vivo* during intraperitoneal glucose tolerance tests (IPGTT), in the presence or absence of the GLP-1 receptor agonist, exendin-4.

Examined in wildtype mice of either sex, acute injection of dorzagliatin or exendin-4 alone exerted only minor effects on glucose tolerance, which did not reach statistical significance (Fig. 5A-D). On the other hand, in wildtype male (but not female) mice, co-injection of dorzagliatin markedly potentiated the action of exendin-4 (Fig. 5A-D).

**Figure 5.**
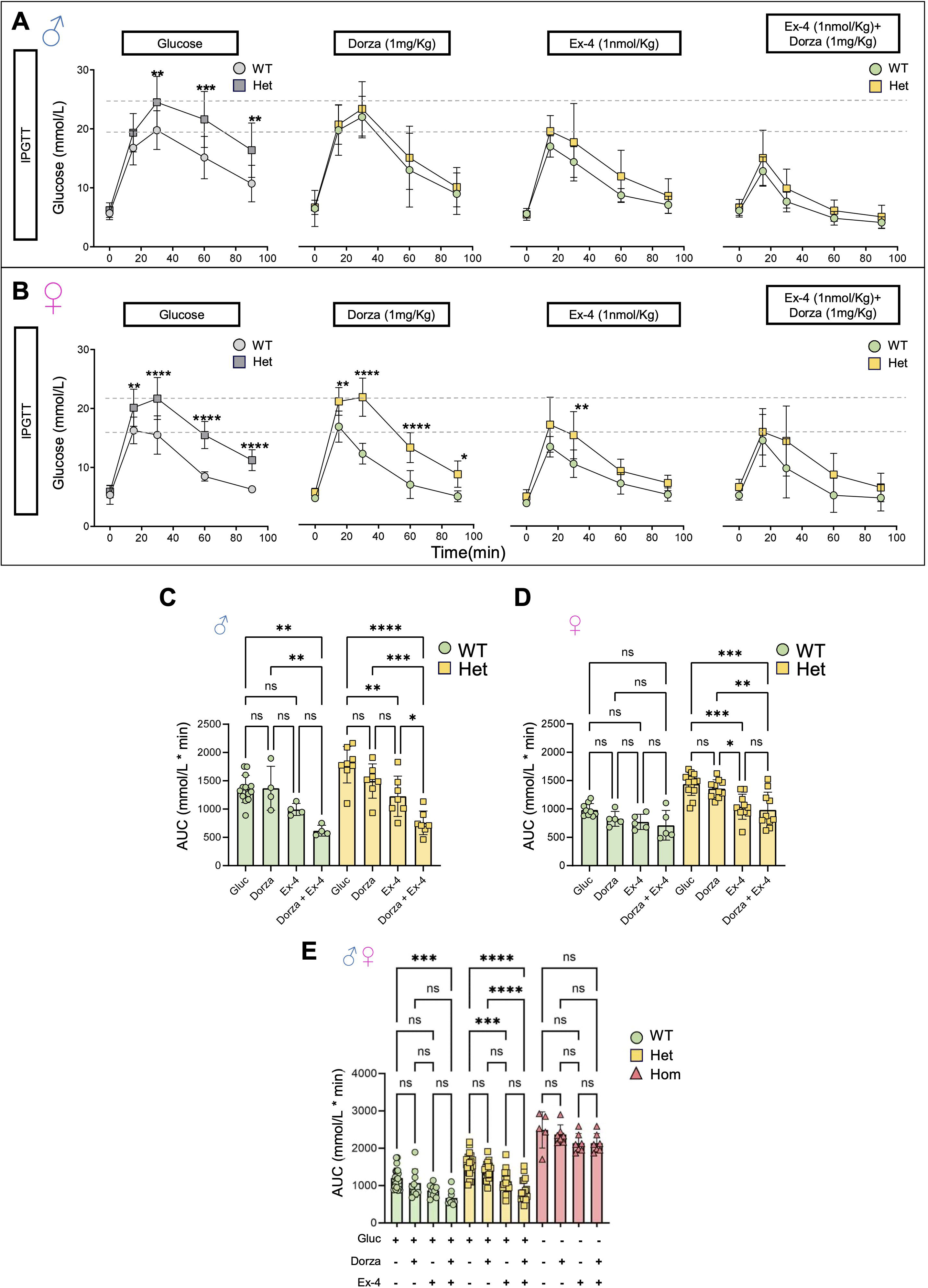
Impact of dorzagliatin and exendin-4 on glucose tolerance across genotypes and sexes. **A-B**) Intraperitoneal glucose tolerance test (IPGTT) with 2 g/kg glucose performed in animals of 8-16w of age for males and females after injection of glucose only or glucose in combination with dorzagliatin (1mg/kg), exendin-4 (1nmol/kg) or dorzagliatin (1mg/kg) + exendin-4 (1nmol/kg) in WT (Gck^+/+^) or heterozygous (Gck^KI/+^) animals. The first panel in grey corresponds to the same data shown in Fig. 3D. Unpaired two-way ANOVA with Šidák’s correction. **C-D)** Glucose AUC quantifications of each IPGTT treatment from male and female mice presented in (A) and (B). Males, *n*=4 WT, 7 Hets for each treatment; Females, *n*=5 WT, 10 Hets for each treatment. Unpaired one-way ANOVA with Tukey’s correction. **E)** Glucose AUC quantifications of each IPGTT treatment from pooled male and female mice presented in (A) and (B), and homozygous (Gck^KI/KI^) mice treated with PBS alone, or PBS and dorzagliatin (1mg/kg), exendin-4 (1nmol/kg) or dorzagliatin (1mg/kg) + exendin-4 (1nmol/kg). Males: *n*=4 WT, *n*=4 Hets, *n*=2 Homo for each treatment, *n*=1-Homo (PBS); Females *n*=7 WT, *n*=7 Hets, *n*=5 Homo for each treatment, *n*=4 Homo (PBS). Unpaired one-way ANOVA with Tukey’s correction. *P ≤ .05, **P ≤ .01, ***P ≤ .001, ****P ≤ .0001 \**P* ≤ .05, \*\**P* ≤ .01, \*\*\**P* ≤ .001, \*\*\*\**P* ≤ .0001.

In male heterozygous Gck^KI/+^ mice, exendin-4 alone, but not dorzagliatin, improved glucose tolerance, whilst the combination of exendin-4 and dorzagliatin profoundly lowered glucose excursions to levels comparable to those in wildtype mice (Fig. 5, A,C). In female Gck^KI/+^ mice, dorzagliatin again exerted no effect when administered alone, whereas exendin-4 markedly improved glucose tolerance. No additional effect was observed of co-injecting dorzagliatin and exendin-4 (Fig. 5 B,D).

Combining data from both sexes, we noted that neither drug, alone or in combination, significantly affected glycemia in homozygous (Gck^KI/KI^) animals (Fig. 5E).

### Effects of incretin and GKA on glucose-stimulated insulin secretion

Explored *in vivo* during IPGTTs, glucose-induced increases in circulating insulin tended to be potentiated by incretin in male, but not female, Gck^+/+^ and Gck^KI/+^ mice (SFig. 6). Since inter-animal variation was large in these experiments, we examined the potential interaction between the GKA and incretin on insulin secretion *ex vivo*. In islets isolated from wildtype male mice, neither dorzagliatin nor exendin-4 significantly affected insulin secretion stimulated by 17 mmol/L glucose (Fig. 6A). In contrast, the combination of dorzagliatin and exendin-4 stimulated hormone release 2.0-2.5-fold. In islets from male Gck^KI/+^ mice (Fig. 6A), dorzagliatin alone tended to increase secretion, whilst exendin-4 alone caused a ∼3-fold increase in hormone release. The effect of incretin was further augmented by dorzagliatin (∼5-fold increase *versus* 17 mmol/L glucose alone). Qualitatively similar results were obtained in female wildtype mice (Fig. 6B). In female Gck^KI/+^ islets, the effects of dorzagliatin or exendin-4, whether administered alone or together, were weaker than those in Gck^KI/+^ males (Fig. 6B *versus* Fig. 6A).

**Figure 6.**
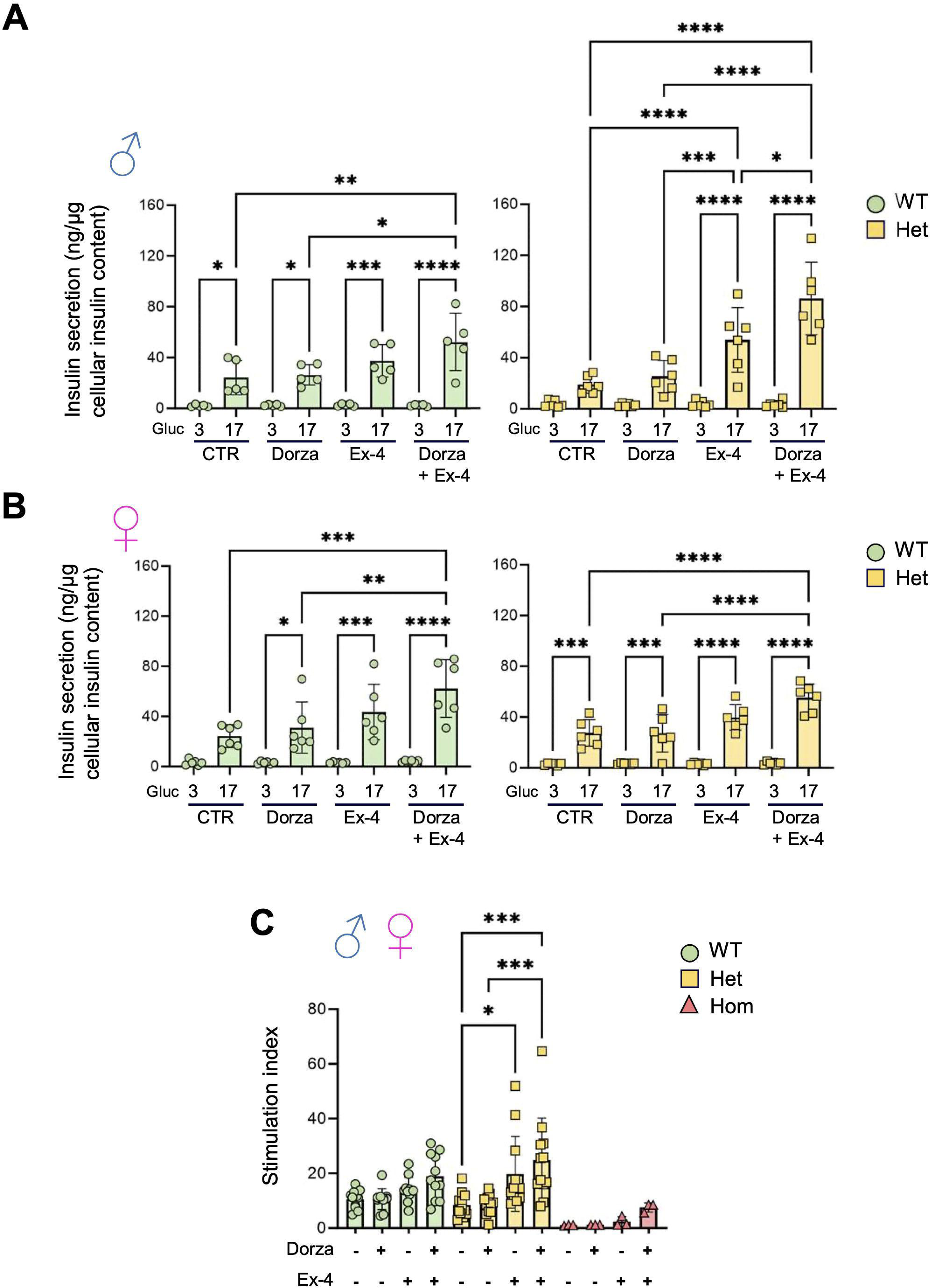
Impact of dorzagliatin and incretin on glucose-stimulated insulin secretion *in vitro*. **A-B**) Insulin secretion *in vitro* for male (A) and female (B) at low (3mmol/L) and high (17mmol/L) glucose concentrations. **C)** Stimulation index, calculated as the ratio of insulin secreted at 17 mmol/L glucose to that at 3 mmol/L glucose, from pooled islets of both sexes across genotypes. One-way ANOVA of Brown-Forsythe. Mean ± SD. *P<0.05, **P<0.01, ***P<0.001, ****P<0.0001.

When data from both sexes were pooled, exendin-4 was found to potentiate glucose-stimulated insulin secretion in Gck^KI/+^ islets, an action further enhanced by dorzagliatin (Fig. 6C). Similar, albeit nonsignificant, tendencies were observed in wildtype and in Gck^KI/KI^ islets (Fig. 6C).

## Discussion

### Novel models of “glucokinase” diabetes

We describe here novel mouse models of *GCK*-MODY and *GCK*-PNDM, and explore the potentiation of glucose-stimulated insulin secretion by a glucokinase activator and incretin in each case.

More than 600 variants are associated with human *GCK*-MODY, with eight identified in *GCK*-PNDM (47). The latter vary in their effects on GK activity or stability, and have broadly proportional clinical impacts (21). Whilst homozygosity for mutants with near-zero activity is usually implicated in PNDM, compound heterozygosity for an inactivating (e.g. (intervening sequence 8 [IVS8] + 2T→G) and hypomorphic (G264S) variant (16) can also drive neonatal diabetes. On the other hand, milder mutations e.g. G223S, L315H, with relative activities of 0.25 and 0.89, respectively (48), lead to *GCK*-MODY. Interestingly in the context of the present study, Osbak *et al* have speculated that milder *GCK* mutations in *GCK*-PNDM e.g. R397L (16; 49) may be more responsive to GKA than those leading to more complete inactivation e.g, T168A (50).

The hypomorphic *Gck* allele described here was originally designed to preserve an intact glucokinase protein, whilst independently expressing, from the same transcript, the fluorescent reporter mCardinal. Instead, an unexpected splicing event eliminated exons 2 and 3 (encoded by the host genome) and led to the fusion of exon 1 to exon 4, derived from the introduced *Gck* cDNA (exons 4-11). The aberrantly-spliced transcript is predicted to produce an inactive protein, lacking much of the ATP– and glucose-binding catalytic domain of the smaller lobe (31). It is also possible that the variant transcript fails to produce any protein product at all. Our Western blotting approach was unable to distinguish between these possibilities, since the product of the aberrantly-spliced variant is unlikely to be detected by the N-terminal targeting antibody used (Results).

In any case, limited correct splicing appears to produce low levels of the active, wildtype protein (10-15% of normal values). Consequently, the Gck^KI^ allele provides a convenient model of inactivating *GCK* mutations that affect the kinetic properties of GK as well as of variants that lead to thermal instability and protein degradation (51).

Enzymatic measurements of GK were not performed since these require beta cell purification to exclude contamination from hexokinases I-III abundant in other cell types (52). Nevertheless, the metabolic features of homozygous *Gck*^KI/Ki^ mice (Fig. 3,5), studies of glucose regulated Ca^2+^ dynamics (Fig. 4; SFig 5) and insulin secretion (Fig.s 3,6) are consistent with near-complete elimination of GK activity from the beta (and other pancreatic endocrine) cells. The persistence of residual glucokinase activity in beta cells likely explains why homozygous Gck^KI/KI^ mice survive into adulthood, in contrast with models of complete *Gck* inactivation (23) (see Introduction). Of note, the metabolic phenotype of heterozygous (Gck^KI/+^) mice was almost indistinguishable from that of previous models involving complete inactivation of a single allele, and thus is compatible with loss of 40-45% of GK activity. Ca^2+^ imaging revealed the expected defects in glucose signaling in both Gck^KI/+^ and Gck^KI/KI^ islets, and the expected rescue of these deficits by dorzagliatin in the former, reflecting the activation of the remaining glucokinase. Whilst the glucokinase activator was largely inactive in male Gck^KI/KI^ islets, as anticipated, 10-30 % of islets displayed responses to dorzagliatin in females (Fig. 4F,G). Interestingly, these findings suggest that correctly spliced, active, GK may be restricted to a subset of Gck^KI/KI^ islets.

Our observation of lowered beta cell mass in homozygous mice aligns with earlier studies in glucokinase-deficient mice (53). Less expected was the increase in alpha cell mass in Gck^KI/KI^ islets, suggesting that GK activity plays a role in suppressing alpha cell expansion, in addition to inhibiting glucagon secretion (54). Extrapolated to humans, these findings raise the intriguing possibility that elevated glucagon levels may aggravate hyperglycemia in *GCK*-PNDM.

We note that, in addition to islets and liver, glucokinase is expressed in several brain nuclei, including the hypothalamus, pituitary and brain stem (33), as well as in intestinal L-cells (55). As such, the phenotypes of Gck^KI/+^ and Gck^KI/KI^ mice may in part reflect actions on these cell types.

### Sex-dependent additive effects of dorzagliatin and incretin on glucose tolerance and insulin secretion

Previous work with earlier-generation GKAs (e.g. the AstraZeneca molecule, GKA50) (56), reported a marked left-shift in the response to glucose in both mouse and human islets. Under the conditions used here, involving high glucose concentrations in both settings, dorzagliatin alone elicited minimal effects either *in vivo* on glucose tolerance (Fig 5 C,D,E), or on glucose-stimulated insulin secretion (Fig. 6A,B) both in wildtype and Gck^KI/+^ mice. Nevertheless, the GKA markedly potentiated the effects of exendin-4 on glucose-stimulated insulin secretion both *in vivo* (Fig. 5C) and *in vitro* (Fig. 6A), an effect most dramatic in heterozygous male mice. The mechanisms underlying this apparent sexual dimorphism in drug responses remain unclear.

### Conclusions

We describe here new mouse models of *GCK*-MODY and *GCK*-PNDM, which may be valuable for future drug screens as well as studies of diabetes complications, including nephropathy. We demonstrate the remarkable efficacy in the former of combining a GKA with a GLP-1R agonist, likely reflecting a requirement for beta cell glucose metabolism for incretin action (28). Our findings suggest the potential therapeutic utility of this drug combination in selected *GCK*-MODY or *GCK*-PNDM patients.

## Supporting information

All Supp Figures

Supp. Tabe 1

SMovie 1

SMovie 2

SMovie 3

SMovie 4

SMovie 5

## Acknowledgements

We thank the CRCHUM Cell Imaging, Animal and Cellular Physiology Facilities for their assistance and Jannick Bonenfant (UdeM) for help with imaging experiments. We are grateful to Drs Khalil Boudaydan, Demetra Rodaros and Thierry Alquier (CRCHUM) for fluorescence measurements in liver and brain sections.

## Author contributions

GAR and MOH conceived the project. GAR designed the studies, supervised the project and wrote the manuscript with input from all authors. SS and LD performed *in vivo* metabolic analyses, and studies of Ca^2+^ dynamics, alongside GO and KD. KB, DR, WT and TA performed histological studies and immunohistochemistry of liver and brain sections. IK, JT, FM and MJH analysed kidney sections with PC, who performed *in vitro* insulin secretion analyses with RM and quantified beta/alpha cell mass. LD, KD, RM and SS performed analysis of alternative splicing, and LD structural modelling with AlphaFold. G.A.R. serves as guarantor of the study.

