## Supplementary material for "Sex-dependent additive effects of dorzagliatin and incretin on insulin secretion in a novel mouse model of *GCK*-MODY": All Supp Figures

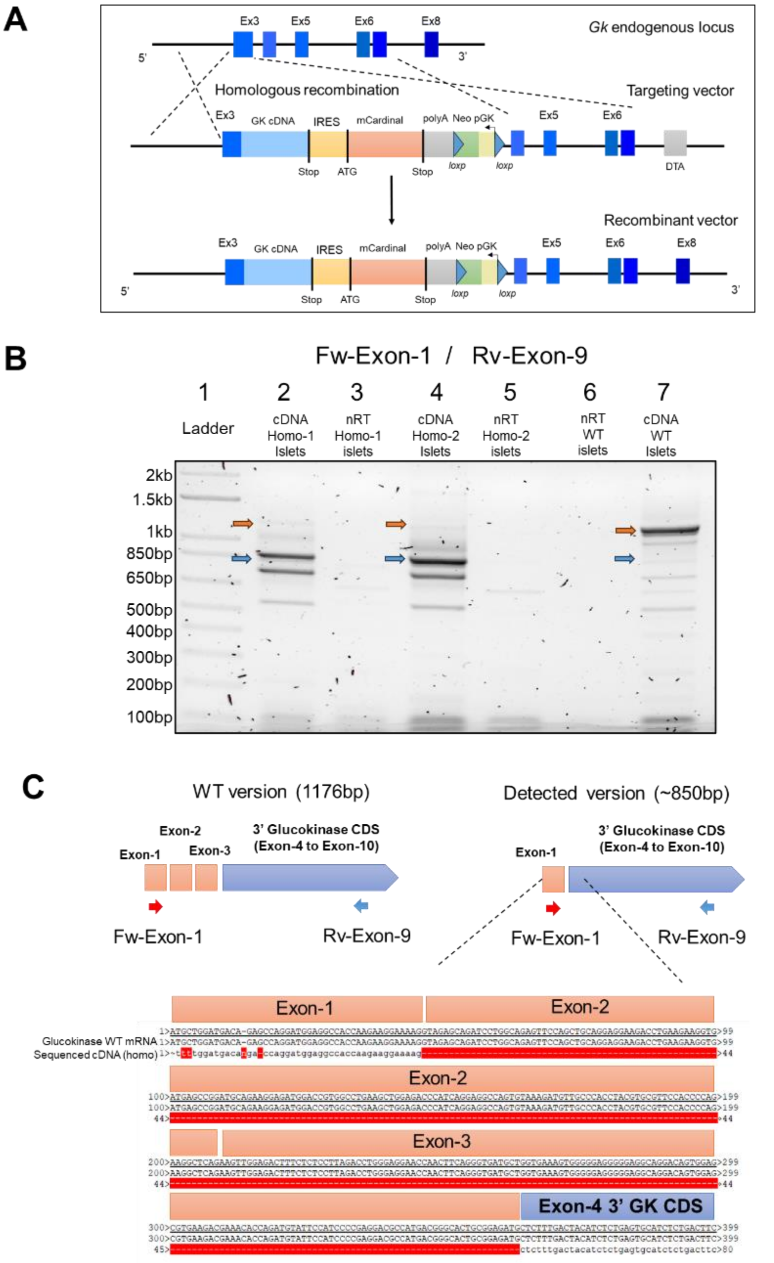


**SFig. 1. Genomic map of the $Gck^{KI}$ mouse allele and cDNA PCR products from $Gck^{KI/KI}$ homozygous and WT cDNA**. **A)** Genomic design from the mouse chromosome 11 targeting the *glucokinase* gene. The 3' *glucokinase* CDS covering the Exon-4 to Exon-10 were directly fused in frame with the endogenous Exon-3 of *glucokinase*, through homologous recombination. **B)** image from a 1% agarose gel after loading the PCR product using the primers to amplify the *glucokinase* mRNA covering Exon-1 to Exon-9. The PCR template was the cDNA of pancreatic islets from $Gck^{KI/KI}$ homozygous males and females pool from 3 animals each and a pool of 3 male WT controls. non-RT (no reverse transcriptase) reactions were added as negative controls. The first lane contains the DNA size ladder. The lane 7, shows the expected band from WT islets. The homozygous yielded 1 faint band of similar molecular size of around 1.17kb (orange arrow). However, the homozygous presented an abundant amplicon of ~850bp. **C)** Sanger sequencing from the ~850bp purified PCR band shown in (A) from $Gck^{KI/KI}$ islet cDNA. The alignment in sequence shows alternative splicing in which Exon-2 and Exon-3 are skipped in comparison to the WT version.

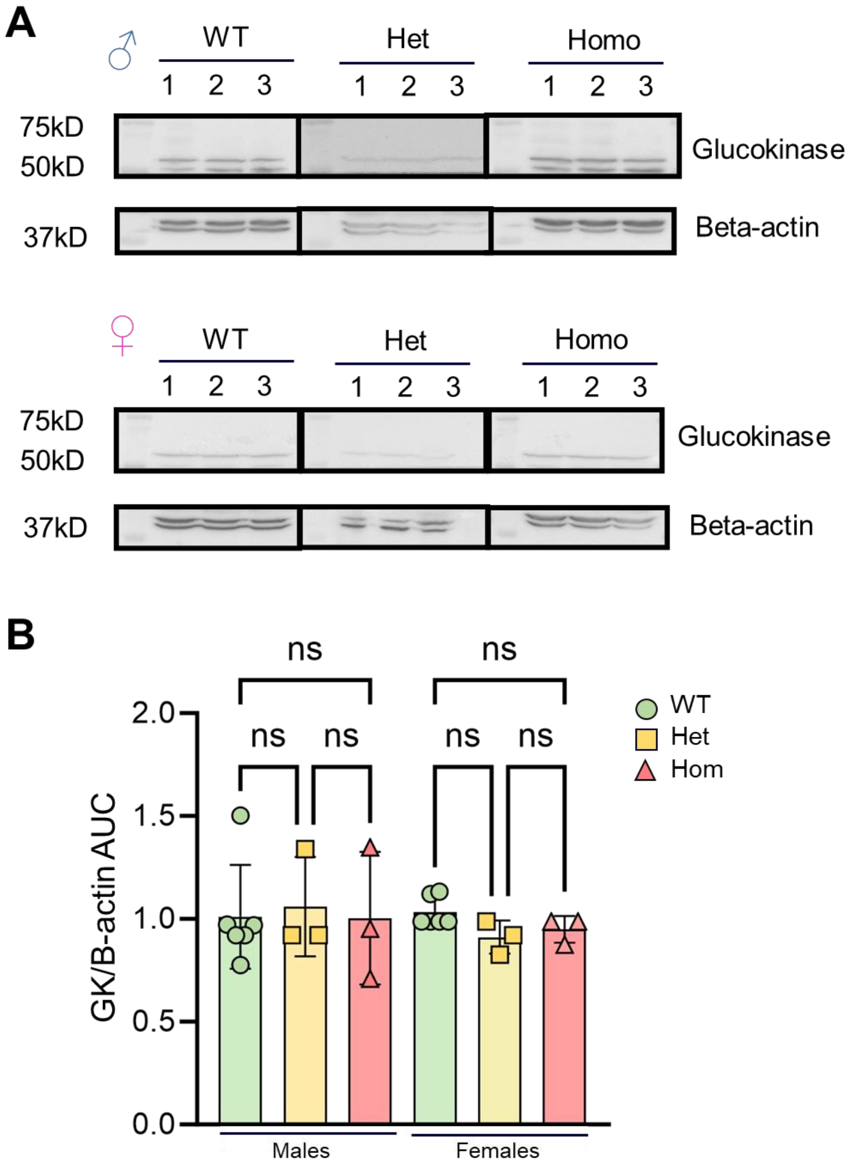


**SFig. 2. Apparent glucokinase immunoreactivity in male and female livers. A)** Western blot images from WT, heterozygous and homozygous males and female liver. A band, presumed to be unrelated to GK, was detected at ~50kDa and B-actin was detected at ~45kDa. **B)** Quantitative analysis of apparent expression levels normalized by B-actin across sex and genotype. No significant differences were found in comparison to WT controls. Each dot represents the quantification of an independent animal. Males: n=6 WT, 3 Hets and 3 Homo. Females: n=6-WT, 3-Hets and 3-Homo. Mice were 8-12w of age. Unpaired one-way ANOVA with Tukey's correction. ns= Not significant.

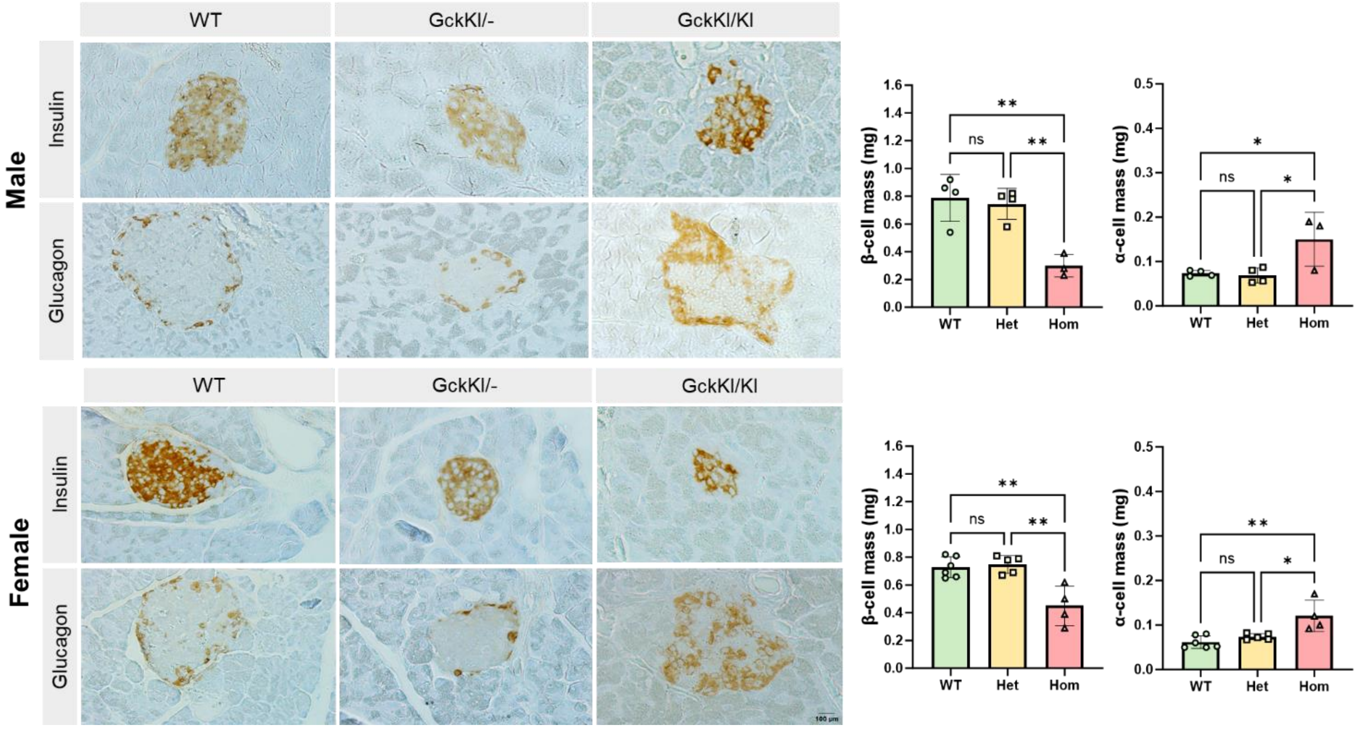


**SFig. 3. Impact of the genotype on beta and alpha cell mass in males and females. A-B)** Representative images and quantitative analysis of beta and alpha cell mass across different genotypes and sexes. One-way ANOVA of Brown-Forsythe. Mean ± SD. *P<0.05, **P<0.01, ***P<0.001.

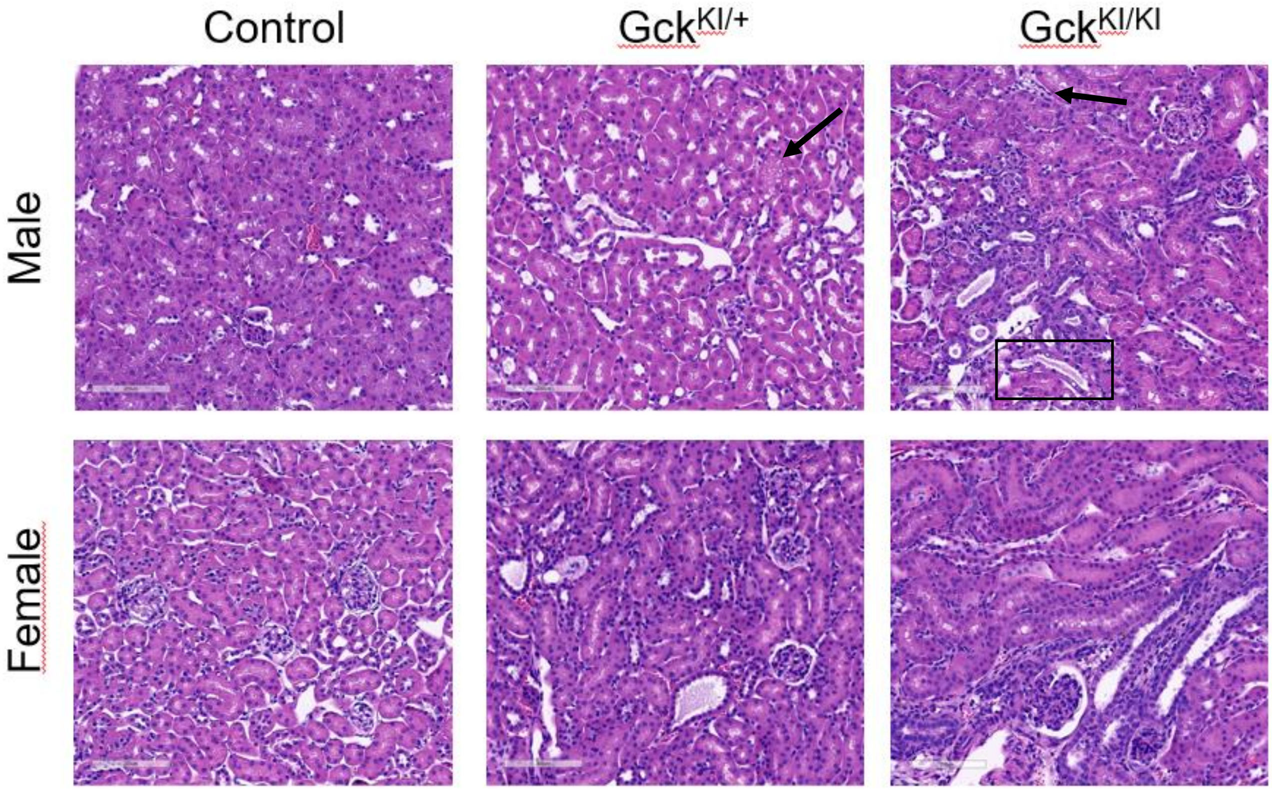


**SFig. 4. Qualitative analysis of H&E-stained renal tissue.** Representative kidney images of male and female mice across the three genotypes. Compared to control mice, heterozygous Gck$^{KI/+}$ mice have numerous small clear vacuoles in tubular epithelial cells (arrow) and mild to moderate tubular damage, whereas the tubules were moderately to severely damaged with frequent epithelial vacuolization (arrow) and chromatin margination (square) in the homozygous Gck$^{KI/KI}$ group. Scale bars: 200µm (magnification x100). See also Supp. Table 2.

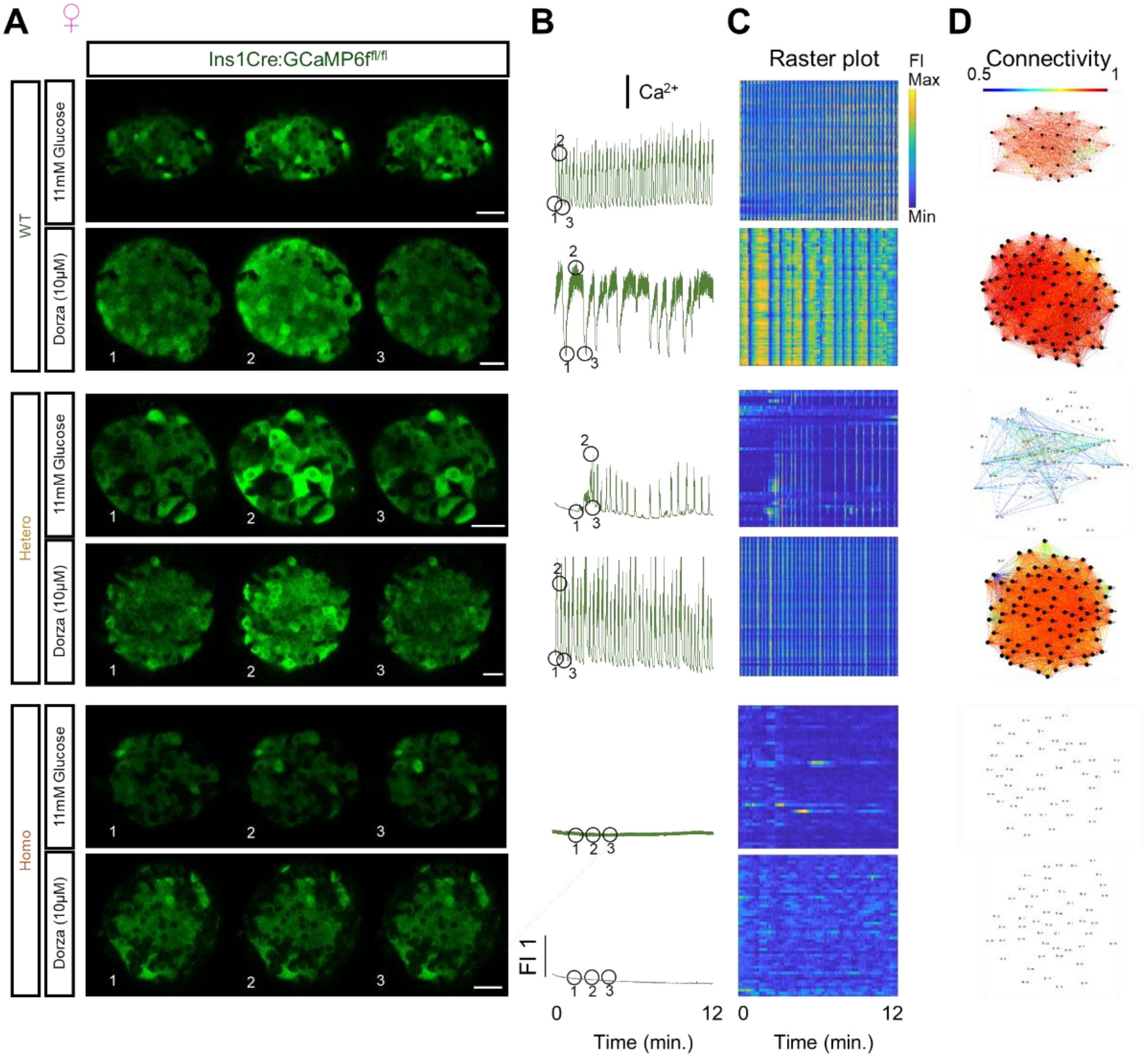


**SFig. 5. Dorzagliatin restores $Ca^{2+}$ dynamics and beta cell connectivity in female $Gck^{KI/+}$ mouse islets. A)** Snapshots from confocal time-lapses recordings of the individual islets at 6 Hz in 11mM glucose with or without 10μM Dorzagliatin between $Gck^{+/+}$and $Gck^{KI/+}$. **C)** Normalized GCaMP6f fluorescence traces from islets shown in (B). **C)** Raster plots show the GCaMP6f signal for individual cells from islets shown in (B). **D)** Visualization of islet connectivity-topography for each genotype and with or without dorzagliatin from islets displayed in (B). The numbers 1, 2, and 3 in the snapshots displayed at (B) indicates the time points corresponding to the islet fluorescent traces showed in (C). Scale bar, 25μM.

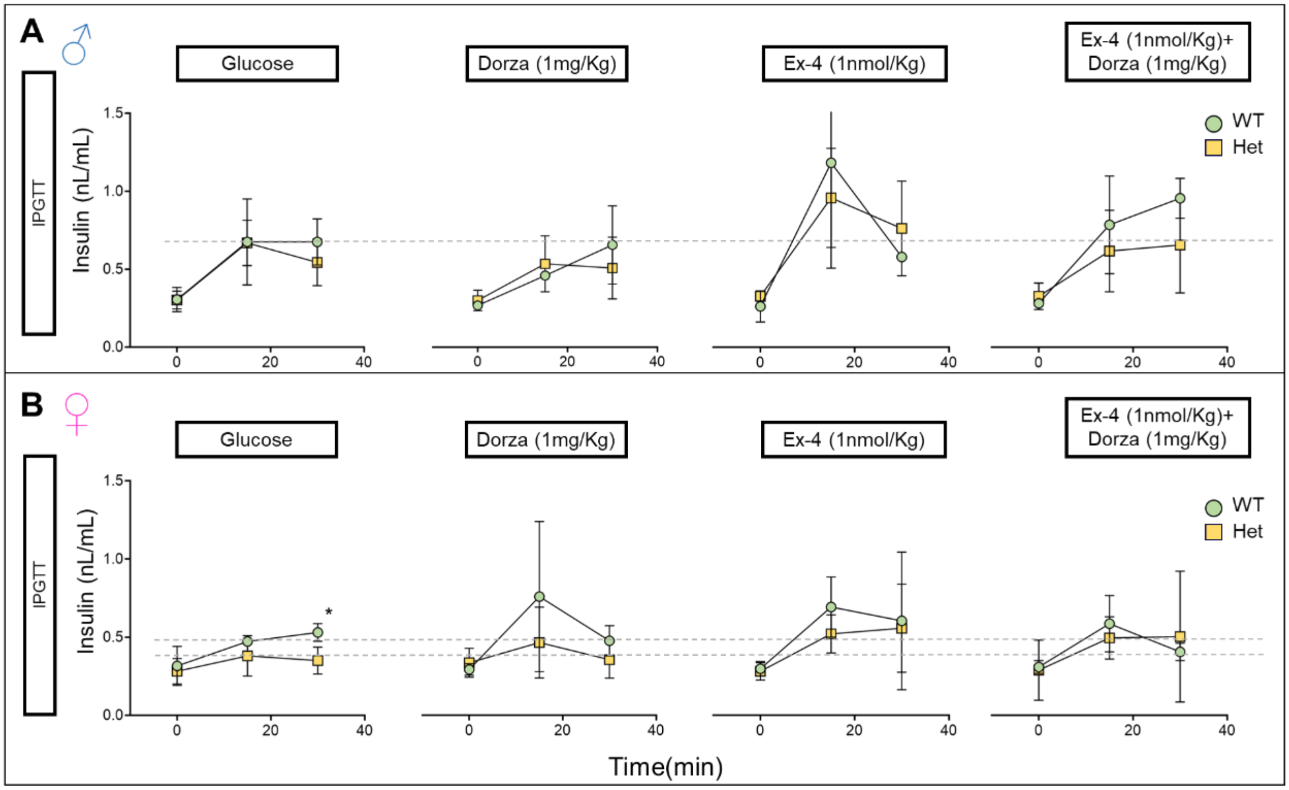


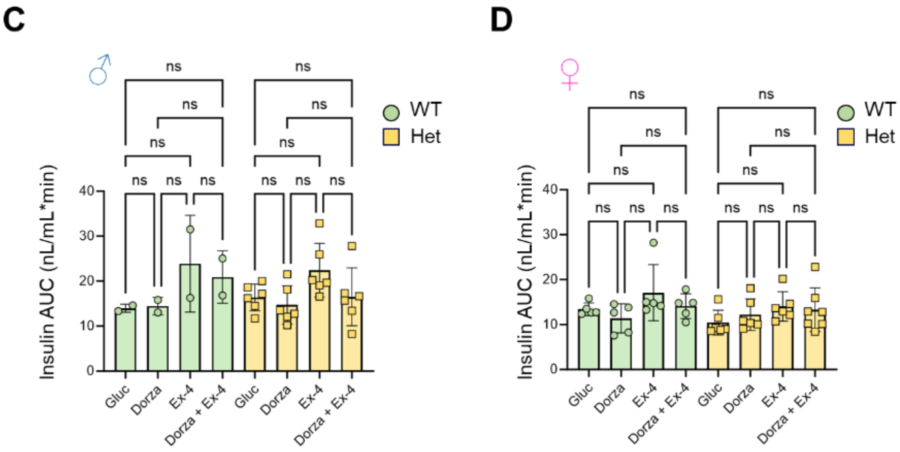


**SFig. 6. Impact of dorzagliatin and Exendin-4 on circulating insulin levels across *Gck* genotypes and sexes. A-B)** Intraperitoneal glucose tolerance test (IPGTT) with 3 g/kg glucose/bw performed at 8-16w of age for males and females with only glucose or in combination with dorzagliatin (1mg/kg), exendin-4 (1nmol/kg) or dorzagliatin (1mg/kg) + exendin-4 (1nmol/kg) in WT or heterozygous background. The graph shows the blood circulatory insulin levels. Unpaired 2-way ANOVA with Šidák's correction. **C-D)** Insulin AUC quantifications of each IPGTT from male and female mice presented at (A) and (B). Males: glucose *n*=2 WT, *n*=6 Het; dorzagliatin *n*=2 WT, *n*=6 Het; exendin-4 *n*=2 WT, *n*=6 Het; and dorzagliatin + exendin-4, *n*=2 WT, *n*=6 Het. Females: dorzagliatin *n*=5 WT, *n*=6 Het; exendin-4 *n*=5 WT, *n*=6 Het; and dorzagliatin + exendin-4, *n*=5 WT, *n*=7 Het. Unpaired one-way ANOVA with Tukey's correction. *$P \leq .05$, ns, not significant.

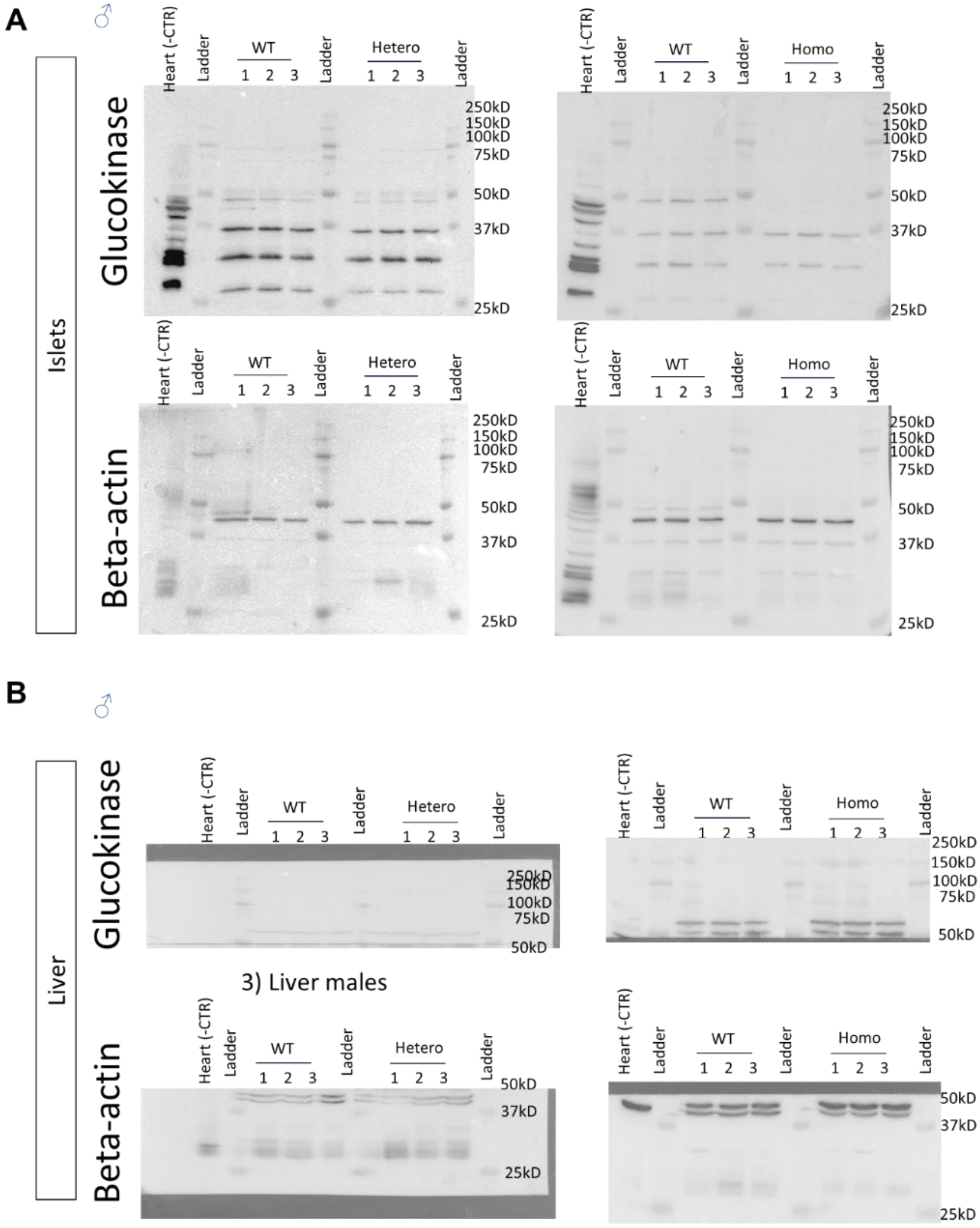


**SFig.8. Whole blots from Glucokinase protein expression levels in male' islet and liver A)** Western blot images from islet protein extracts of WT, heterozygous and homozygous males. The Glucokinase was detected at ~45kDa and B-actin was detected at ~40kDa. At least 3 unspecific bands were also detected by the Gk-primary antibody at around ~35kDa, ~30kDa and ~28kDa in islet protein extracts. **B)** Western blot images from liver protein extracts of WT, heterozygous and homozygous males. The Glucokinase was detected at ~50kDa and B-actin was detected at ~40kDa. At least one unspecific band was detected by the Gk-primary antibody at around ~50kDa in liver protein extracts.

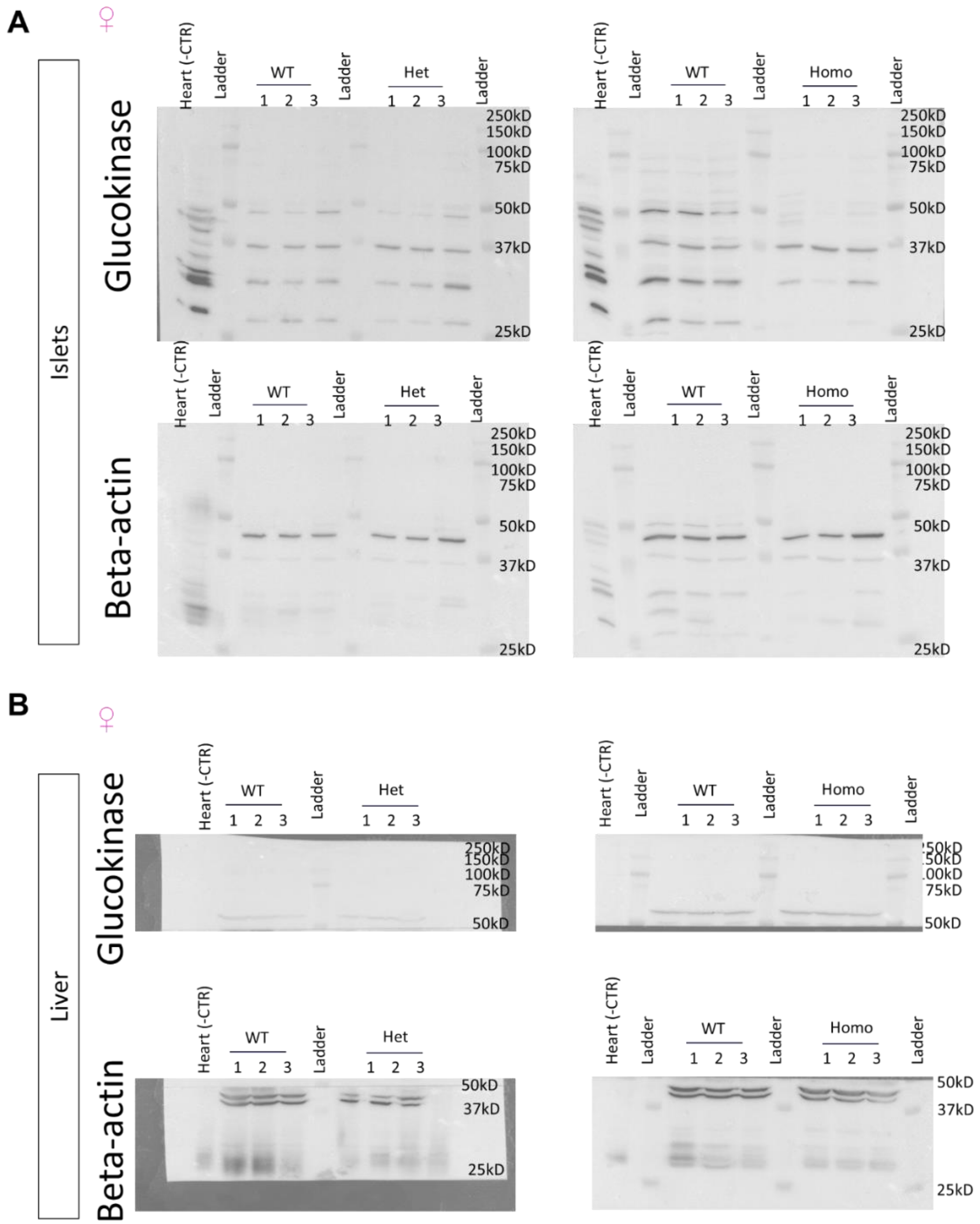


**SFig. 9. Whole Western-blots from Glucokinase protein expression levels in female' islet and liver A)** Western blot images from islet protein extracts of WT, heterozygous and homozygous females. The Glucokinase was detected at ~45kDa and B-actin was detected at ~40kDa. At least three unspecific bands were also detected by the Gk-primary antibody at around ~35kDa, ~30kDa and ~28kDa in islet protein extracts. **B)** Western blot images from liver protein extracts of WT, heterozygous and homozygous males. Glucokinase was detected at ~50kDa and B-actin was detected at ~40kDa.

### Videos.

**Movie S1. AI 3D models.** 3D model animation of AlphaFold predicted Glucokinase of mouse WT and the predicted miss-spliced isoform.

**Movie S2. Confocal calcium imaging across male genotypes.** Calcium imaging of pancreatic islets expressing GCaMP6f specifically in beta-cells was performed in male islets from WT, heterozygous and homozygous. The oscillatory changes of GCaMP fluorescence signal report the glucose-stimulated influx of calcium during the imaging session. The video was recorded at 6Hz with stimulatory glucose (11mM).

**Movie S3. Confocal calcium imaging across male genotypes upon 10μM dorzagliatin treatment.** Calcium imaging of pancreatic islets expressing GCaMP6f specifically in beta-cells was performed in male islets from WT, heterozygous and homozygous. The islets were treated with 10μM dorzagliatin and stimulatory glucose (11mM). The video was taken at 6Hz.

**Movie S4. Confocal calcium imaging across female genotypes.** Calcium imaging of pancreatic islets expressing GCaMP6f specifically in β-cells was performed in female islets from WT, heterozygous and homozygous. The oscillatory changes of GCaMP fluorescence signal reports the glucose-stimulated influx of calcium during the imaging session. The video was recorded at 6Hz with stimulatory glucose (11mM).

**Movie S5. Confocal calcium imaging across female genotypes upon 10μM dorzagliatin treatment.** Calcium imaging of pancreatic islets expressing GCaMP6f specifically in β-cells was performed in male islets from WT, heterozygous and homozygous. The islets were treated with 10μM dorzagliatin and stimulatory glucose (11mM). The video was taken at 6Hz.
