## Supplementary material for "Sex-dependent additive effects of dorzagliatin and incretin on insulin secretion in a novel mouse model of *GCK*-MODY": Supp. Tabe 1

**Supplementary Table 1.** Primers used to amplify cDNA from Gck^KI/KI^ transcript.

| Forward primers |  |
| --- | --- |
| Fw-Exon-1 | GGATGACAGAGCCAGGATGGAGGCCACCAAGAAGG |
| Fw-8-mCard | GTACGGGAGCAAGACCTTCATCAACCACAC |
| Fw-9-mCard | TACTGCGACCTCCCTAGCAAACTGGGGCAC |
| Fw-9-mCard | TACTGCGACCTCCCTAGCAAACTGGGGCAC |
| Fw-Exon10 | CACCCGAGCTTCAAGGAGCGGTTTCACGCC |
| Fw-Exon-3 | GTTGGAGACTTTCTCTCCTTAGACCTGGGAGGAA |
| Fw-Exon-7 | GATGAGAGCTCAGTGAACCCCGGTCAGCAG |
| Fw-IRES | TTGCAGGCAGCGGAACCCCCCACCTGGCGA |
| Reverse primers |  |
| Rv-8-mCard | GTGTGGTTGATGAAGGTCTTGCTCCCGTAC |
| Rv-9-mCard | GTGCCCCAGTTTGCTAGGGAGGTCGCAGTA |
| Rv-Exon10 | GGCGTGAAACCGCTCCTTGAAGCTCGGGTG |
| Rv-Exon-3 | TTCCTCCCAGGTCTAAGGAGAGAAAGTCTCCAAC |
| Rv-Exon-7 | CTGCTGACCGGGGTTCACTGAGCTCTCATC |
| Rv-Exon-9 | GCGATTTATGACCCCCGCTAGTCCTGCTG |
| Rv-Exon10 | GGCGTGAAACCGCTCCTTGAAGCTCGGGTG |
| Rv-IRES | TCGCCAGGTGGGGGGTTCCGCTGCCTGCAA |
| Rv-8-mCard | GTGTGGTTGATGAAGGTCTTGCTCCCGTAC |
| Rv-9-mCard | GTGCCCCAGTTTGCTAGGGAGGTCGCAGTA |
| Rv-PolyA | TGGCCCTGGAAGTTGCCACTCCAGTGCCCACC |
| Rv-PolyA-2 | CCTTCTATAATATTATGGGGTGGAGGGGGG |
