## Supplementary material for "Sex-dependent additive effects of dorzagliatin and incretin on insulin secretion in a novel mouse model of *GCK*-MODY": Supp. Table 2

Supplementary Table 2. Renal Histology. See Text and Supp Fig. 4

| Sex | [illegible]notype | Observations | |
|---|---|---|---|
| | | Tubular compartment | Glomerular compartment |
| Male | WT | Mild tubular damage | Absence of damage |
| Male | WT | Mild tubular damage and few vacuolized epithelial cells | Absence of damage |
| Female | WT | Absence of tubular damage | Absence of damage |
| Female | WT | Absence of tubular damage and rare chromatin margination | Absence of damage |
| Female | WT | Mild tubular damage and frequent vacuolization of epithelial cells | Absence of damage |
| Female | WT | Mild tubular damage | Absence of damage |
| Female | WT | Absence of tubular damage | Absence of damage |
| Male | Het | Mild tubular damage, frequent chromatin margination and abundant vacuolization of epithelial cells | Presence of a periglomerular fibrosis |
| Male | Het | Moderate tubular damage | Focal segmental glomerulosclerosis |
| Male | Het | Mild tubular damage | Absence of damage |
| Female | Het | Mild tubular damage and few chromatin margination | Absence of damage |
| Female | Het | Moderate tubular damage | Absence of damage |
| Female | Het | Mild tubular damage and slight vacuolization of epithelial cells | Absence of damage |
| Male | Homo | Moderate tubular damage and vacuolization of epithelial cells | Absence of damage |
| Male | Homo | Severe tubular damage with chromatin margination and isometric vacuolization of epithelial cells | Absence of damage |
| Male | Homo | Moderate tubular damage and abundant chromatin margination | Presence of a sclerotic glomerule |
| Female | Homo | Moderate to severe tubular damage and abundant chromatin margination | Absence of damage |
